# Re-charging your fats: Charmm36 parameters for neutral lipids triacylglycerol and diacylglycerol

**DOI:** 10.1101/2021.09.29.462351

**Authors:** Pablo Campomanes, Janak Prabhu, Valeria Zoni, Stefano Vanni

## Abstract

Neutral lipids (NLs) are an abundant class of cellular lipids. They are characterized by the total lack of charged chemical groups in their structure, and, as a consequence, they play a major role in intracellular lipid storage. NLs that carry a glycerol backbone, such as triacylglycerols (TGs) and diacylglycerols (DGs), are also involved in the biosynthetic pathway of cellular phospholipids, and they have recently been the subject of numerous structural investigations by means of atomistic molecular dynamics (MD) simulations. However, conflicting results on the physicochemical behavior of NLs were observed depending on the nature of the atomistic force field used. Here, we show that current phospholipid-derived CHARMM36 parameters for DGs and TGs cannot reproduce adequately interfacial properties of these NLs, due to excessive hydrophilicity at the glycerol-ester region. By following a CHARMM36-consistent parameterization strategy, we develop new parameters for both TGs and DGs that are compatible with both cutoff-based and Particle Mesh Ewald (PME) schemes for the treatment of Lennard Jones interactions. We show that our new parameters can reproduce interfacial properties of NLs and their behavior in more complex lipid assemblies. We discuss the implications of our findings in the context of intracellular lipid storage and NLs cellular activity.

---

Neutral lipids (NLs), generally defined as naturally occurring hydrophobic molecules that lack any charged chemical group, are an important class of cellular lipids(Gurr and James 1980). As a consequence of their extreme hydrophobic character, they are immiscible in water, and, above their transition temperature, they do not self-assemble in lamellar phases, such as lipid bilayers, but rather behave as pure liquids. They are thus characterized by classical properties of liquids, including density, surface tension (ST) with the gas phase, and interfacial tension (IT) with water.

Amongst NLs, most of the attention has focused on cholesterol, both because of its abundance in the cell and its implications in countless cellular processes and diseases. As a consequence, cholesterol is also the most studied NL from a structural perspective. To this extent, computational investigations of cholesterol using molecular dynamics (MD) simulations, the method of choice to investigate lipid assemblies at the structural level(Enkavi et al. 2019; Marrink et al. 2019), date back to the 90s’(Edholm and Nyberg 1992; Robinson et al. 1995; Tu et al. 1998).

Recently, however, other NLs involved in phospholipid synthesis and degradation, and most notably triacylglycerol (TG) and diacylglycerol (DG), have received increasing attention at the structural level(Hall et al. 2008; Khandelia et al. 2010; Vuorela et al. 2010; Ollila et al. 2012; Vamparys et al. 2013; M’barek et al. 2017; Tascini et al. 2018; Campomanes et al. 2019; Zoni et al. 2021a, b, 2019; Chorlay et al. 2019), in large part due to ever-increasing focus on cellular lipid storage and related metabolic diseases(Welte 2015). To this extent, both fully atomistic (all-atom, AA) and coarse-grain (CG) MD simulations of TG- and DG-containing systems have been reported, using different force fields(Hall et al. 2008; Khandelia et al. 2010; Ollila et al. 2012; Bacle et al. 2017; Prévost et al. 2018; Campomanes et al. 2019; Olarte et al. 2020; Kim and Swanson 2020; Caillon et al. 2020; Prasanna et al. 2021; Zoni et al. 2021a, b; Kim et al. 2021; Kim and Voth 2021).

Historically, the first AA parameters for NLs were prepared based on the OPLS-compatible Berger parameterization strategy(Hall et al. 2008), and they have been successfully used to investigate several structural aspects of high-density and low-density lipoproteins(Vuorela et al. 2010; Ollila et al. 2012), lipid droplets-like surfaces(Bacle et al. 2017) and oil-water interfaces(Tascini et al. 2018). More recently, new parameters compatible with the widely successful CHARMM36 (C36) force fields for lipids have been used to investigate the properties of lipid assemblies containing TGs, in the absence or presence of accompanying proteins(Olarte et al. 2020; Kim and Swanson 2020; Prasanna et al. 2021). These parameters, C36-s (C36-standard) from now on, are derived from those developed for phosphatidyl-choline (PC) lipids by simply replacing the charged phosphate and choline groups with one additional acyl chain in position *sn*-*3*.

Quite remarkably, however, the two parameter sets (Berger *vs* C36-s) produce contrasting results on key physicochemical properties of TG assemblies. Most notably, the two models suggest very different hydration properties for TG, with the C36-s model showing ~10 times more water molecules in bulk liquid TG(Kim and Swanson 2020) with respect to simulations carried out with Berger parameters(Bacle et al. 2017). This is even more confounding when considering that the reported IT of triolein (TOG) with water is very similar (31 ± 2 mN/m for Berger(Ollila et al. 2012), 29.7 ± 1.7 mN/m for C36-s(Kim and Swanson 2020)) for the two models and, in both cases, close to the reported experimental values (29.2(Couallier et al. 2018) and 32 mN/m(Mitsche et al. 2010)).

From a biological perspective, the simulations carried out with the more hydrophilic C36-s set of parameters have led to a model where TG molecules (named “SURF-TG”) can reside at the surface of lipid bilayers, adopting phospholipid-like conformations and acting as monolayer lipids(Kim and Swanson 2020). This interpretation has potentially profound implications for what pertains to both integral and peripheral proteins interactions with lipid droplets (LDs), as well as for the mechanism of TG degradation by lipases(Olarte et al. 2020; Kim et al. 2021).

To investigate the incongruity between the two parameter sets, and since the results with the Berger model have been validated by a number of independent research groups(Ollila et al. 2012; Bacle et al. 2017; Tascini et al. 2018), we opted to start by computing the IT of TOG via MD simulations on a flat TOG-water interface using the C36-s parameterization (see Supporting Information, SI, for further details). As can be appreciated in Figure 1, the IT estimated from the analysis of the trajectory obtained when the C36-s model was employed for TOG diverges from the expected range of experimental measurements (29-32 mN/m), rather plateauing to a value^1^ of 17.3 ± 2.0 mN/m after a relatively long equilibration period. Of note, a similar discrepancy between the experimental IT value and that obtained with the C36-s model has been reported in a recent manuscript(Kim and Voth 2021). Interestingly, we observed that this decrease in IT is coupled to a linear increase in the hydration of the TOG core (Figure 1), which could explain the long simulation time required to equilibrate this system (approximately 400 ns). At this stage, the amount of water that penetrates into the oil core stabilizes to some extent, and the IT profile displays the expected convergence fingerprint, which allows estimating an IT average value for this system. This observation provides a likely explanation for the difference between our result (17.3 mN/m) and that previously reported(Kim and Swanson 2020) (29.7 mN/m) since, in that study, the IT for TOG was computed from a short 50 ns run.

**Figure 1.**
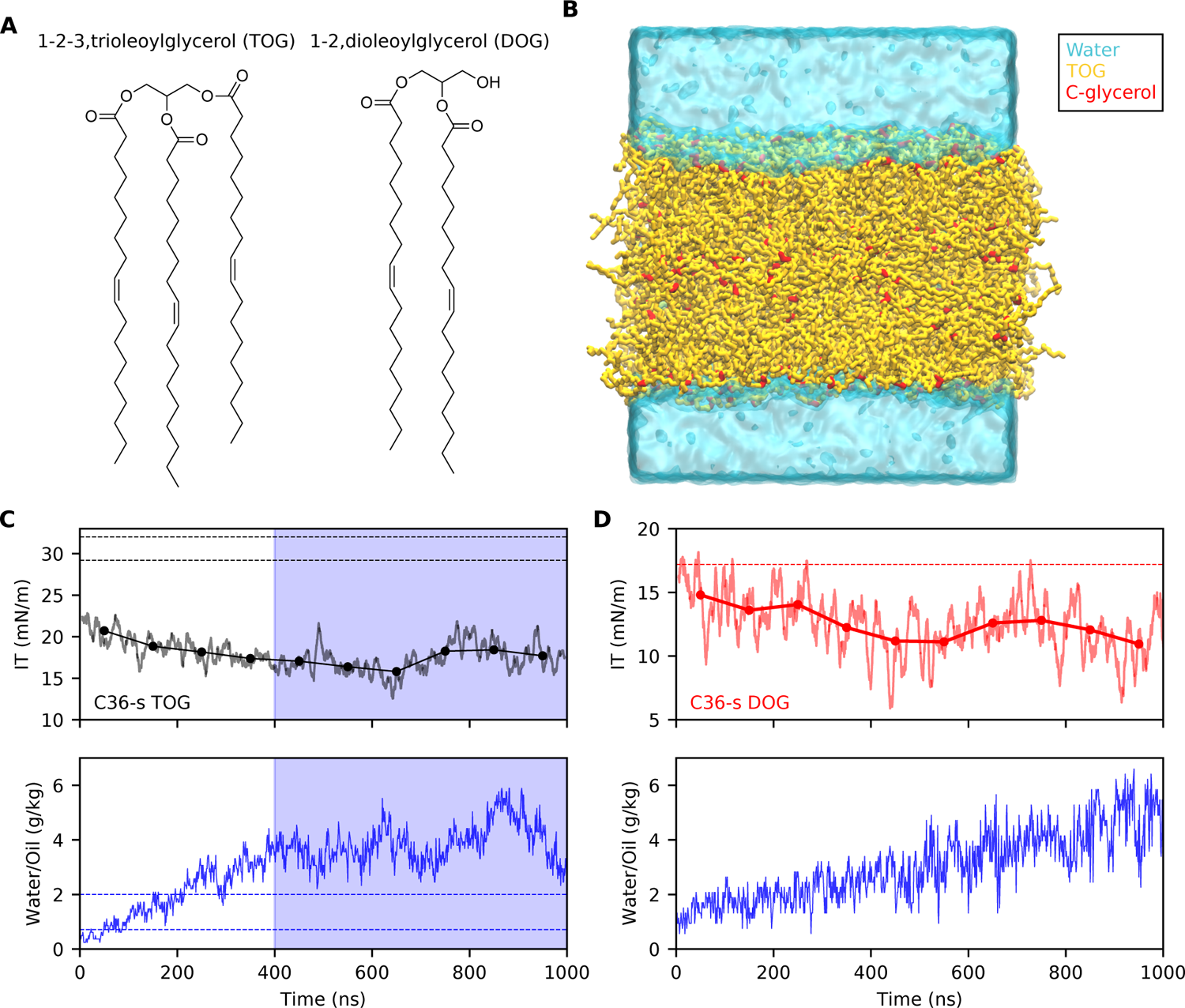
Interfacial tension of triolein (TOG) and dioleoyl-glycerol (DOG) computed from simulations carried out using the C36-s model. (A) Chemical structures of TOG and DOG. (B) Setup for interfacial tension (IT) calculations. Water is represented in cyan, TOG in yellow, and the carbons of the glycerol group (C1, C2, C3) in red. (C) IT values for TOG as running averages (solid lines) and blocking averages (mean values every 100 ns) (top) and water content inside the oil core (bottom). The horizontal dashed lines represent the experimental values reported for both IT (top, black) and water content (bottom, blue). The blue shaded region contains the part of the trajectory used to calculate the mean IT value for TOG, when water content shows convergence. (D) IT values for DOG as running averages (solid lines) and blocking averages (mean values every 100 ns) (top) and water content inside the oil core (bottom). The horizontal dashed line represents the experimental value reported for DOG IT (top, red).

We next investigated whether a similar criticality was present in another NL, dioleoylglycerol (DOG). DOG is the natural precursor of both TOG and cellular phospholipids, and, unlike TOG, is generally considered as a *bona fide* bilayer lipid, even though its presence above 20-30% molar concentration in phospholipid bilayers promotes the formation of non-lamellar structures(Das and Rand 1986). Indeed, for DOG we observed a similar behavior to that of TOG (Figure 1); in this case, even if the simulation seems to still lack convergence because water is still penetrating into the oil core after a long 1 μs-run, the IT at the DOG water interface calculated using the C36-s model significantly deviates from that experimentally measured (17.2 mN/m(Nakajima 2004)).

To solve this issue, and especially considering that the C36 force field is widely used in MD simulations of lipid systems, we next investigated the origin of the inability of the C36-s model to reproduce this key experimental parameter. We noticed that, during the development of the C36 parameters for lipids, the charges on the ester groups of PC lipids, from which those on TG and DG molecules derive, were explicitly increased with respect to the QM-derived charges in order to reproduce the correct hydration of PC lipids(Klauda et al. 2010). Thus, in order to minimally affect C36 parameters that have shown very good agreement with experimental data(Klauda et al. 2010), we opted to reparameterize exclusively the charges on the glycerol and ester groups using CHARMM36 compatible strategies(MacKerell et al. 1998; Klauda et al. 2010). Those present in the acyl chains and the alcohol group (in the case of DOG) were not altered and therefore kept at their original C36 values.

In short, we initially got a conformationally-consistent set of CM5 atomic charges(Marenich et al. 2012) for TOG that, as dictated by CHARMM parameterization philosophy, was determined from a population analysis of quantum wave functions determined by implicitly taking into account water-TOG interactions. Then, these initial set of charges was fine-tuned following a regularized gradient-based iterative procedure similar to that previously outlined(Yu et al. 2021a) (see SI for further details on the parameterization protocol). With the new parameters obtained after running two cycles of the above-described iterative procedure (Figure 2A), the IT values for TOG and DOG converged to 31.2 ± 1.4 and 19.1 ± 1.4 mN/m, respectively (Figure 2B). These are in close agreement with those experimentally measured. It must be remarked that this optimal set of parameters, C36-cutoff (C36-c), was developed considering a cutoff-based treatment of the Lennard-Jones (LJ) interactions and can therefore be used in combination with the existent 12Å-cutoff C36 force fields for proteins(Best et al. 2012; Huang et al. 2017) and phospholipids(Klauda et al. 2010; Yu et al. 2021a).

**Figure 2.**
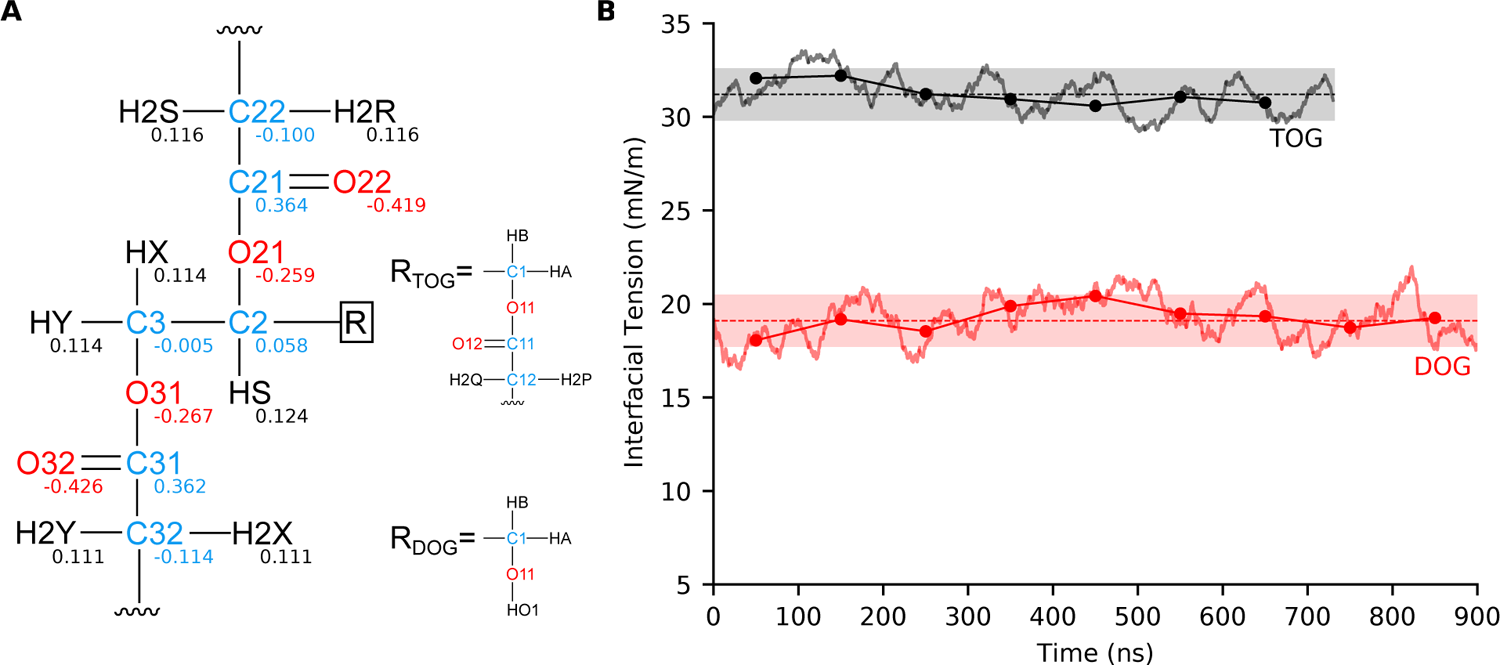
Interfacial tension of triolein (TOG) and dioleoyl-glycerol (DOG) computed from simulations carried out using the C36-c model. (A) Fine-tuned set of charges for TOG/DOG. For TOG, the charges of atoms in chain 1 are identical to those shown for chain 3. (B) Interfacial tension (IT) values for TOG and DOG as running averages (solid lines) and blocking averages (mean values every 100 ns) (dotted solid lines). Only the equilibrated part of the trajectories is shown. Mean values (dashed horizontal lines) and their corresponding 95% confidence intervals extracted from the simulations are also displayed.

We next decided to assess the ability of the C36-c model to reproduce other experimental properties in molecules containing the same glycerol-ester skeleton found in TGs and DGs. To this end, we investigated whether, using the C36-c set of parameters, it was possible to accurately estimate TOG density, and the density and surface tension of a set of molecules (Figure 3A) for which these two properties have been experimentally measured(Yaws 2014). Of note, while densities are in relatively good agreement with the experimentally reported values (showing relative errors below 5%), surface tensions are not well-reproduced by the C36-c model and present relative errors in the 15-23% range. However, this is a well-known issue of the C36 lipid model(Yu et al. 2021b, a) that, as shown for several alkanes and oils(Leonard et al. 2018), originates from the cutoff-based treatment of LJ interactions.

**Figure 3.**
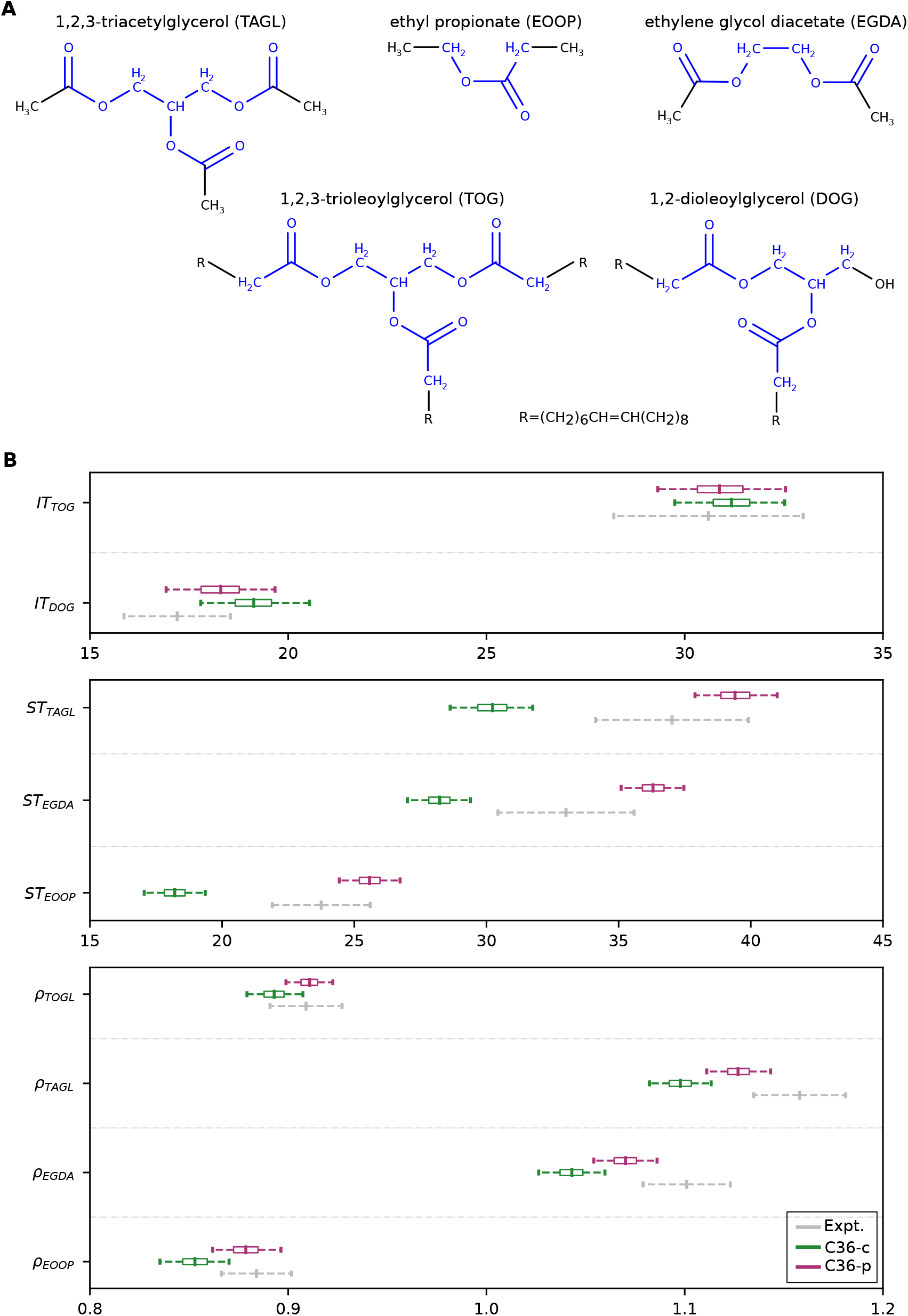
Parameterization approach for CHARMM36 force field using Lennard-Jones-PME (C36-p). (A) Set of molecules used as training set to carry out the C36-p parameterization. The atoms for which charges were optimized are colored in blue. (B) Densities (ρ, in g/cm^3^), surface tensions (ST, in mN/m), and interfacial tensions (IT, in mN/m) experimentally measured (gray) and computed using the C36-c (green) and C36-p (magenta) sets of parameters. Mean values and 95% confidence intervals are shown. The boxes represent the first and third quartiles of the distributions sampled during the dynamics. Confidence intervals for the experimental measurements were estimated as described in SI.

Fortunately, a solution to this problem has been recently proposed via the use of a PME-like treatment of the Lennard-Jones interactions in C36 (LJ-PME)(Yu et al. 2021b, a). We thus opted to also develop C36 LJ-PME compatible parameters for TGs and DGs. To do so, we used a thermodynamic reweighting procedure similar to that described above, but targeting not only TOG and DOG IT, but also TOG density as well as density and surface tension of the molecules displayed in Figure 3A. By doing so, we also expect a larger model transferability. We used the optimal set of charges of the C36-c model to initiate the protocol, which was then fine-tuned as described above but activating now the usage of LJ-PME in our simulations. After a single iteration, we obtained a new set of charges, C36-PME (C36-p) (Figure S1), that was able to reproduce fairly well all the properties targeted (Figure 3B). In particular, the mean relative errors for densities and ST were about 1% and 8%, respectively. Moreover, as shown in Figure 3B, this new set of charges jointly led to TOG and DOG IT values (30.9 ± 1.6 and 18.3 ± 1.4 mN/m, respectively) that are in very good agreement with the reported experimental measurements.

As an independent additional validation, we next estimated the amount of water dispersed in the oil core in the case of TOG (for which this property has been experimentally measured(Ragni et al. 2012)) from the simulations performed with the C36-c and C36-p models (Figure 4). As mentioned above, it appears that there is a correlation between the IT values and the water content inside the oil core. This correlation is not unexpected as both properties are related to the hydrophilicity of the oil molecules. In particular, when the C36-c and C36-p models were used, mean values of 0.98 and 1.01 g/kg water/oil, respectively, were found for the water-holding capacity of TOG. These computed values are in good agreement with the experimentally reported range of 0.8-2 g/kg water/oil for TG samples(Ragni et al. 2012) thus giving some initial support to the adequacy and improved accuracy of the parameter sets developed in this study.

**Figure 4.**
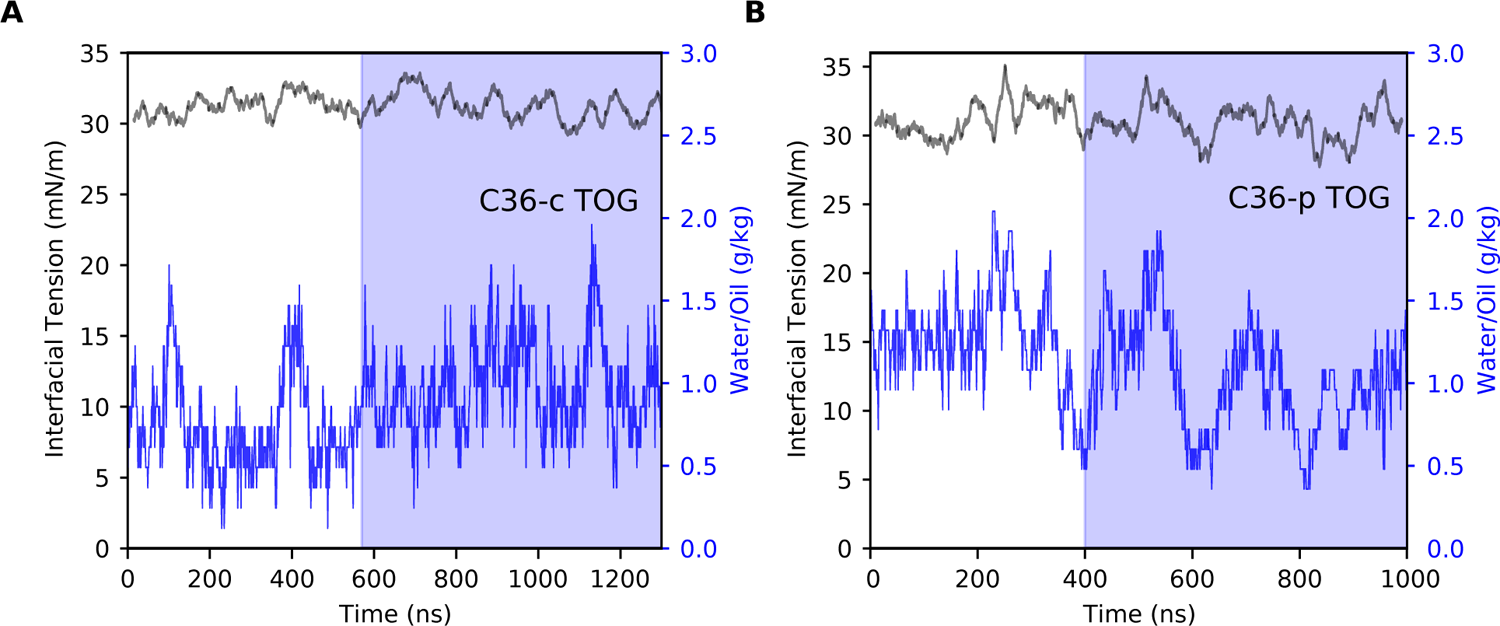
Interfacial tension (black) and water content (blue) of triolein (TOG) computed from simulations carried out with (A) the C36-c model and (B) the C36-p model. The blue shaded region contains the part of the trajectory used to calculate the mean IT value for TOG, when water content shows convergence.

Finally, to further validate the accuracy of the parameter sets developed in this work (C36-c and C36-p), we investigated the behaviour of DOG and TOG in the presence of bilayer-forming lipids, as this is the most common setup for the study of physiologically relevant properties of NLs. To do so, we estimated the flip-flop energy barrier of physiologically-abundant monounsaturated TG and DG molecules (TOG and DOG) in palmitoyl-oleyl (PO)-PC lipid bilayers using either C36-c or C36-p for the NLs in combination with the corresponding CHARMM force fields (C36 or C36-LJPME, respectively) for the phospholipids (Figure 5). The potentials of mean force (PMF) shown in Figure 5 were computed using Boltzmann inversion from four independent MD simulations using different initial configurations for the various replicas (see SI for further details). The obtained average values for the flip-flop energy barriers amount to about 1 kcal/mol for TOG and 3.5 kcal/mol for DOG. The latter is in good agreement with that experimentally estimated for DG molecules(Hamilton et al. 1991), which further supports the strategy employed for the parameterization and the accuracy of the charges here developed for the glycerol-ester groups.

**Figure 5.**
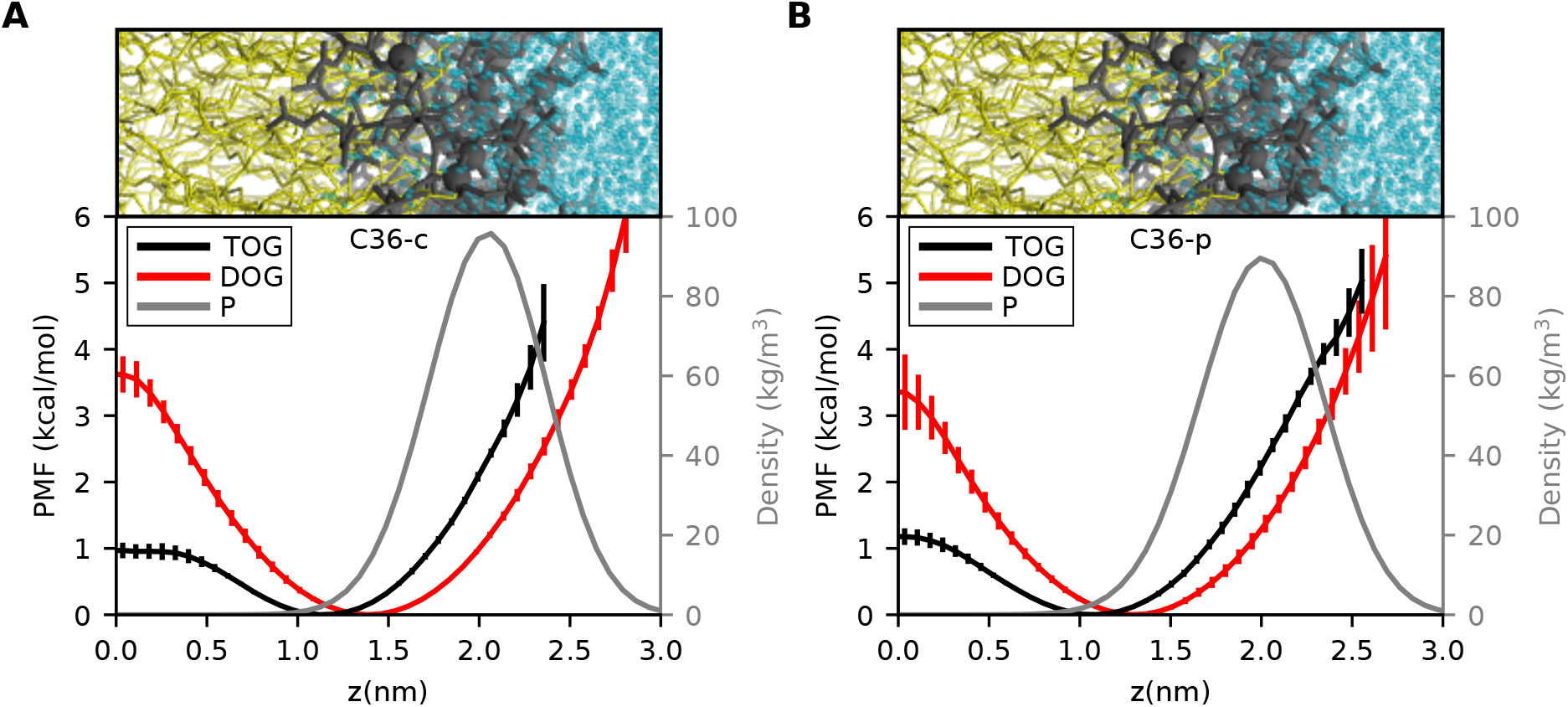
Potentials of Mean Force (PMF) calculated for TOG (black) and DOG (red) in POPC bilayers. (A) Results for simulations obtained using the new C36-c parameters for TOG and DOG. (B) Results from simulations with the new C36-p parameters for TOG and DOG. The PMF curves are obtained as an average over four replicas. The associated uncertainties (standard deviations) are also shown together with the estimated density profile for the phosphate group (P) of POPC (gray line), which was computed as an average between both TOG/POPC and DOG/POPC simulations.

Most notably, our results allow to evaluate the potential biological implications of a (in)correct treatment of the hydrophobic/hydrophilic balance in MD simulations of NLs. Our data indicate that the glycerol moiety of TG molecules is most abundant in the hydrophobic interior of the bilayer, albeit not at the interface between the two monolayers bur rather 1 nm above it (Figure 5). This confirms previous models suggesting that the presence of TG molecules in bilayers can alter the arrangement of bilayer phospholipids(Bacle et al. 2017; Kim and Swanson 2020; Kim et al. 2021). On the other hand, the energetic cost to move from one monolayer to the other is very low (1 kcal/mol), and significantly lower than the cost of reaching the bilayer surface (~2.5 kcal/mol), defined as the average level of the phosphate groups of the phospholipids (Figure 5). Hence, TG flip-flop between monolayers is extremely fast, thus excluding potential monolayer activity of TG molecules that could lead to any sort of TG-induced asymmetric behaviour, such as membrane bending(Kim and Voth 2021) or LD budding(Chorlay et al. 2019), in the presence of symmetric lipid bilayers.

DG, on the other hand, displays a behaviour that is intermediate between that of TG and that of phospholipids(Kornberg and Mcconnell 1971) (Figure 5). As such, it should not be simply thought of as a “classical” glycerolipid, with its polar head pointing towards water and its acyl chains embedded in the hydrophobic core of the lipid bilayer, but rather as a more complex, multi-faceted molecule with a physico-chemical complexity that mirrors its biological one(Campomanes et al. 2019).

In conclusion, we identified significant shortcomings in the current C36 force field parameters for TG and DG molecules. These issues are the origin of reported discrepancies in the description of oil/water and oil/phospholipid/water interfaces using MD simulations. Here, we provide new parameters compatible with the C36 force fields for proteins and phospholipids based on a minimal parameterization strategy that focuses on reducing the excessively high point charges (and their resulting polarization) on the ester and glycerol groups of the molecules. Our new models provide excellent agreement with key physicochemical properties of NLs, and are fully compatible with cutoff-based and PME schemes for the treatment of LJ interactions. We foresee that the parameters developed in this work will be of help to address the ever-increasing interest in the structural role of NLs in modulating key physiological processes related to cell homeostasis and related metabolic diseases.

## Supporting Material

Supporting material includes a description of the methods used in this work, two figures, and one table

## Acknowledgments

This work was supported by the Swiss National Science Foundation (grant #PP00P3_163966). This project has received funding from the European Research Council (ERC) under the European Union’s Horizon 2020 research and innovation programme (Grant agreement No. 803952). This work was supported by grants from the Swiss National Supercomputing Centre (CSCS) under project ID s980 and s1030.

## Author Contributions

The manuscript was written through contributions of all authors. All authors have given approval to the final version of the manuscript.

## Footnotes

1 Unless otherwise stated, from now on, uncertainties will be reported as 95% confidence intervals.

