## Supplemental Information for "Re-charging your fats: Charmm36 parameters for neutral lipids triacylglycerol and diacylglycerol"

**Supplemental Methods**

***Systems setup***

Suitable starting configurations to compute DOG (1,2-dioleoyl-sn-glycerol) and TOG (trioleoylglycerol) interfacial tensions (IT) were obtained by backmapping coarse grained (CG) equilibrated snapshots. These snapshots were extracted from CG MD simulations that used force fields for TOG and DOG developed in previous studies(Bacle et al. 2017; Campomanes et al. 2019). These "oil in water" structures contained an oil core composed by either 512 DOG or 432 TOG molecules immersed in water, and led to equilibrated periodic boxes with sizes of about 85x85x125 $Å^3$ and 105x105x120 $Å^3$, respectively.

This solvated TOG equilibrated structure, upon removal of the water molecules, was also used to initiate the simulations required to compute the oil density. On the other hand, starting random configurations to calculate density and surface tension (ST) for other molecules in the training set (see Figure 3 in the main text) were built using PACKMOL(Martínez et al. 2009). Every system was composed by 512 identical molecules.

The systems used to estimate the DOG and TOG flip-flop rates in POPC bilayers were built according to the following procedure. We prepared four replicas for each of the systems (POPC/DOG and POPC/TOG). Two of them were fully generated using CHARMM-GUI(Jo et al. 2008, 2009) by randomly inserting an equal amount of oil molecules (DOG or TOG) in each of the POPC leaflets. For the other two, we used CHARMM-GUI to initially build initial pure POPC systems; their corresponding leaflets were then separated along the z-direction (perpendicular to the bilayer-water interface) to sandwich a layer of neutral lipids (DOG or TOG) in between. All the POPC/oil systems were composed of 800 lipids. The molar concentration of lipids were 96:4 for POPC:TOG and 90:10 POPC:DOG, and the lipid:water ratio of all the systems was set to 1:50.

***Parameterization protocol***

In order to develop new parameters for TOG and DOG, our working hypothesis was that the topologies commonly used for these molecules, taken from standard repositories(Jo et al. 2008; Kim and Swanson 2020), present atomic point charges on their ester and glycerol groups that are inaccurate and the unique factor responsible for an excessive hydration near this region (see Figure 1 in the main text). Therefore, to maximize the compatibility with the existent CHARMM36 (C36) force fields for phospholipids(Klauda et al. 2010; Yu et al. 2021), we decided to employ a minimal parameterization strategy according to which only the atomic charges on the ester/glycerol skeleton of TOG and DOG were optimized whereas the rest of parameters, particularly the charges on the acyl chains, were kept at their original values. To keep compatibility with the existent C36 schemes to treat the LJ interparticle interactions, cutoff-based and PME approaches, we developed two different set of parameters: C36-c and C36-p, respectively. Their development followed a workflow similar to that previously employed to generate optimal parameters for phospholipids within the C36/LJ-PME framework(Yu et al. 2021). As described below in further detail, besides a different treatment of the LJ interactions, some other distinct choices were taken to build these two models. Concretely, different sets of initial charges and of training targets were employed.

All classical MD simulations in this study were performed under periodic boundary conditions using the version 2020.4 of GROMACS(Pronk et al. 2013; Páll et al. 2015; Abraham et al. 2015) and shared some common settings: i) the long-range electrostatic atomic interactions were taken into account by means of the particle mesh Ewald (PME) algorithm(Essmann et al. 1995) with a Fourier grid space of 0.12 nm and a real space cutoff of 1.2 nm; ii) all bonds involving hydrogen atoms were constrained using the LINCS and SETTLE algorithms(Miyamoto and Kollman 1992; Hess 2008), thus allowing the usage of an integration time step of 2 fs; iii) the TIP3P model(Jorgensen et al. 1983) was employed to describe water molecules in all lipid/water simulations.

*C36-c (C36-cutoff) model*

To get an initial set of atomic charges (fine-tuned at a later stage) for this model, we first sampled the conformational space of a TOG molecule in vacuum by means of an 80 ps-long MD run. From this simulation, which was performed with a semi-empirical method (AM1)(Dewar et al. 1985), we extracted 200 frames, and for each of them we performed a single point energy calculation at the HF/6-31G(d) theory level(Hehre et al. 1986) using the conductor-like polarizable continuum model (CPCM)(Barone and Cossi 1998) to implicitly mimic a water environment. All these electronic structure calculations were performed using the ORCA package(Neese et al. 2020). This protocol allowed us to obtain conformational energies and wave functions at a theory level higher than AM1 while taking into account the effect of water interactions in the determination of the atomic charges, as dictated by CHARMM parameterization philosophy. The above-mentioned wave functions were used to compute CM5 atomic charges(Marenich et al. 2012) for every configuration extracted from the dynamics via a subsequent population analysis and, lastly, these charges were then averaged to get the initial set to be optimized.

The optimization of these CM5 charges was performed using a gradient-based iterative procedure similar to that previously described(Yu et al. 2021). The gradient on the hyper-surface defined by these parameters (atomic charges) was estimated via thermodynamic reweighting(Zwanzig 1954). To carry out this optimization and obtain a set of charges compatible with the glycerol-ester groups present in both triglycerides (TG) and diacylglycerols (DG), DOG and TOG interfacial tensions were used as target properties. We estimated a mean value and a 95% confidence interval for TOG IT from the three independent experimental studies published in the literature(Mitsche et al. 2010; Couallier et al. 2018; Needham et al. 2019). An analogous relative uncertainty (8%) was used to estimate the 95% confidence interval for DOG. These confidence intervals were employed to define the weights for the training targets; we define them as inversely proportional to their corresponding uncertainties. A regularization strategy that restrained the final set of atomic charges to stay as close as possible to the original CM5 set was applied to avoid overfitting. To this end, we used regularization weights that ensured a maximum change of 0.005e on any partial charge during the optimization cycles. At the end of every optimization cycle, mean values and confidence intervals were extracted from the distributions sampled during the dynamics. The iterative protocol was run until enough overlap was observed between the corresponding confidence intervals (experimental data *vs*. simulations).

All MD simulations required for the parameterization were performed in the NPT ensemble. Constant temperature (298.15 K) and pressure (1 atm) were imposed by coupling the system to a stochastic velocity rescaling thermostat(Bussi et al. 2007) with a coupling time constant of 1 ps and a Parrinello-Rahman barostat(Parrinello and Rahman 1981) with a coupling time constant of 5 ps, respectively. The van der Waals interactions were truncated using a cutoff value of 1.2 nm and a standard smoothing function for the tail region (1.0-1.2 nm). Because of the correlation observed in our simulations between water content inside the oil core and IT, we monitored water penetration inside the oil core along the dynamics and used it as a measure of equilibration (see Figures 4 and S2). After relatively long equilibration periods, production runs of at least 500 ns were carried out to collect enough statistics and get accurate estimates for the IT of the different oils (DOG and TOG). The surface tension was computed from the diagonal values of the pressure tensor ($P_{xx}$, $P_{yy}$, and $P_{zz}$), using the Kirkwood-Irving method(Irving and Kirkwood 1950), as follows:

$$\gamma = \frac{L}{2}\langle P_{zz} - \frac{P_{xx} + P_{yy}}{2}\rangle \quad (1)$$

where $L$ represents the box length along z and <…> denotes ensemble average.

*C36-p (C36-PME) model*

To build this model, the C36-c optimal set of atomic charges was employed to initiate the optimization protocol. These charges were fine-tuned according to a procedure similar to that described above for the C36-c model (i.e., via thermodynamic reweighting) but, in this case, the training set was expanded and various properties of compounds containing the glycerol/ester group in their molecular skeleton were selected as training targets (Table S1). To estimate confidence intervals for the experimental densities, we took the experimental data reported in the literature for TOG(Rowe et al. 2009; Elvers 2017) and acted as described above in the case of IT. For surface tensions, we used relative uncertainties of 8% in analogy as those employed for IT experimental values (Table S1). The weights for the training targets had to be scaled in this case since the order of magnitude of the training targets was quite different. The scaling factors used to solve this issue are collected in Table S1. As before, regularization weights were chosen to control the change on the atomic charges and keep them as close as possible to the original set. In this case, the regularization weights were selected to allow a maximum change of 0.02e on any partial charge at every optimization step. At the end of every optimization cycle, mean values and confidence intervals were extracted from the distributions sampled during the dynamics. The iterative protocol was run until enough overlap was observed between the corresponding confidence intervals (experimental data *vs*. simulations). The optimal set of atomic charges for this C36-p model is shown in Figure S1.

Pressure control varied depending on the property to be computed from the simulations: i) isotropic (in all dimensions) for densities; ii) semi-isotropic (xy coupled together, z independently) for IT. On the other hand, no pressure control was applied in the simulations performed to calculate ST; the NVT ensemble was used in this scenario. Constant temperature and pressure (when applicable) were imposed using the same algorithms and parameters described in the previous section (C36-c model). After equilibration (determined, in the case of IT, by a constant average value of the water content inside the oil core), production runs of 25 ns, 100 ns, and not shorter than 500 ns were carried out to calculate density, ST, and IT, respectively. ST and IT were obtained using equation 1 above. In all MD simulations required to build the C36-p model, a real-space cutoff of 1.0 nm was employed to treat the van der Waals interactions at short-range, whereas the recently developed LJ-PME algorithm(Leonard et al. 2018) was used to treat all interparticle interactions at larger distances (long-range).

### *Flip-flop energy barriers for DOG and TOG*

All MD simulations carried out for these POPC/oil/water systems were performed in the NPT ensemble using semi-isotropic pressure control and analogous settings to those described above for the cutoff and PME-like treatment of the LJ interactions in C36. The lateral density profiles for the phosphorus atom type (P) of POPC and carbon (C2) of the ester group of the oils (DOG and TOG) were calculated using the *gromacs density* tool. The number of bin slices for calculating these profiles was set to 120. The potentials of mean force (PMF) were computed by Boltzmann inverting the corresponding density profiles, using the C2 atom as reference. The simulations were carried out for 400-800 ns, until the estimated energy barriers converged. The final part of the trajectories for the different replicas (200-300 ns) was then used for analyses. The final values for the PMF are reported as average and standard deviation over four replicas.

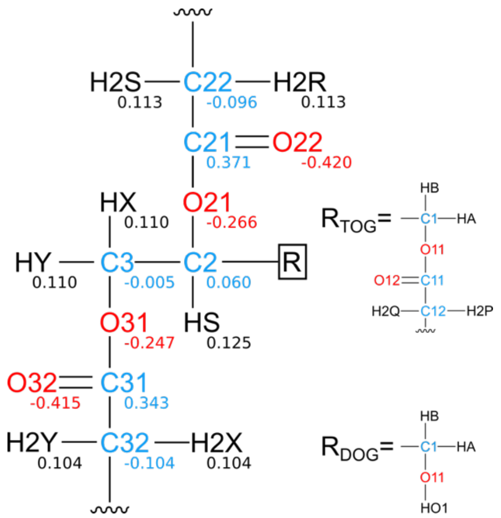


**Figure S1**. C36-p fine-tuned set of charges for TOG/DOG. For TOG, the charges of atoms in chain 1 are identical to those shown for chain 3.

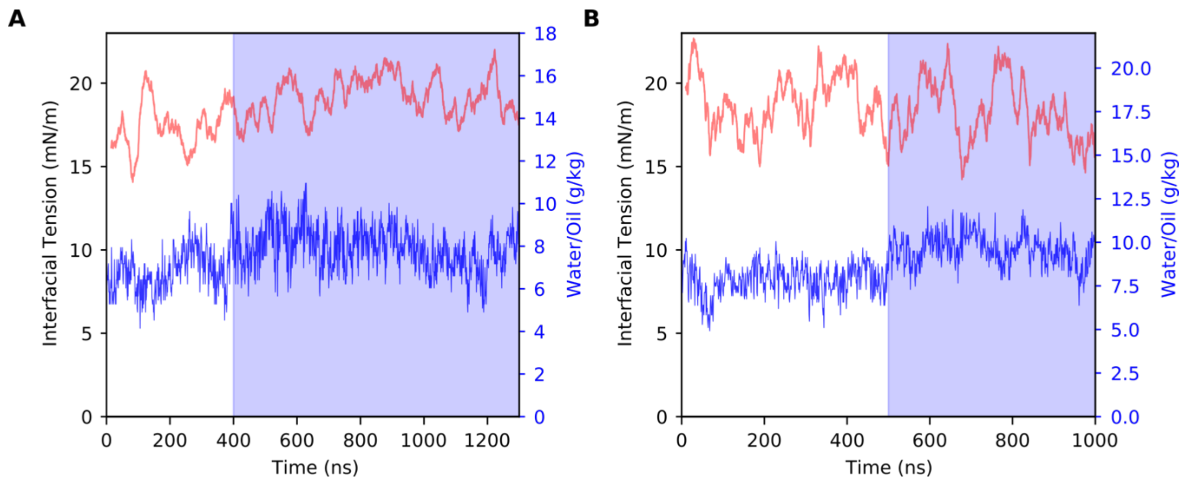


**Figure S2**. Interfacial tension (red) and water content (blue) of DOG computed from simulations carried out with (A) the C36-c model and (B) the C36-p model. The blue shaded region contains the part of the trajectory used to calculate the mean IT value for TOG, when water content shows convergence.

**Table S1**. Training target data (mean values and estimated 95% confidence intervals, CI). Densities are given in g/cm$^3$, while surface tensions at the oil/air interface (ST) and interfacial tensions with water (IT) are given in mN/m. The scaling factors used to make sure that all training targets had a similar order of magnitude during the iterative optimization protocol are also shown. The chemical structures of all systems are displayed in Figure 3A (main text).

| ***System*** | ***Property*** | ***Mean value*** | ***95% CI*** | ***Scaling factor*** |
|---|---|---|---|---|
| **TOG** | IT | 30.6 | 2.4 | 0.040 |
| **DOG** | IT | 17.2 | 1.4 | 0.040 |
| **EOOP** | ST | 23.8 | 1.9 | 0.035 |
| **EGDA** | ST | 33.0 | 2.6 | 0.035 |
| **TAGL** | ST | 37.0 | 3.0 | 0.035 |
| **EOOP** | density | 0.884 | 0.018 | 1.000 |
| **EGDA** | density | 1.101 | 0.022 | 1.000 |
| **TAGL** | density | 1.158 | 0.023 | 1.000 |
| **TOG** | density | 0.909 | 0.018 | 1.000 |
